# A model of tuberculosis clustering in low incidence countries reveals more on-going transmission in the United Kingdom than the Netherlands between 2010 and 2015

**DOI:** 10.1101/639260

**Authors:** Ellen Brooks-Pollock, Leon Danon, Hester Korthals Altes, Jennifer A. Davidson, Andrew M. T. Pollock, Dick van Soolingen, Colin Campbell, Maeve K. Lalor

## Abstract

Tuberculosis (TB) is a major public health threat, including in low TB incidence countries, through a combination of imported infection and onward transmission. Using data from the Enhanced Tuberculosis Surveillance system in the United Kingdom (UK) and the Netherlands (NL) Tuberculosis Register, we developed a mathematical model of TB importation and transmission in low-incidence settings. We apply this method to compare the effective reproduction number for TB, the contribution of importation and the role of superspreaders in the UK and the NL. We estimate that on average, between 2010 and 2015, a TB case generated 0·41(0·30,0·60) secondary cases in the UK, and 0·24(0·14,0·48) secondary cases in the NL. A majority of cases did not generate any secondary cases. Recent transmission accounted for 26% (21%,36%) of UK cases and 19%(11%,31%) of NL cases. We predict that reducing UK transmission rates to those observed in NL would result in 538(266,818) fewer cases annually in the UK. This methodology reveals common transmission mechanisms across two low-incidence countries and can be applied to other settings. Control policies aimed at limiting spread have a role to play in eliminating TB in low incidence countries.

---

Tuberculosis (TB) is a chronic infectious disease and a major global public health threat. In 2017, 10.0 million people developed TB and 1.3 million people died from TB worldwide(1). In many low TB incidence countries, a high proportion of cases occur in persons born abroad, and control measures such as migrant screening have been introduced to limit imported infection and reduce treatment costs(2,3). However, it is often not known whether foreign-born individuals were exposed to TB before or immigration (4,5), which affects the impact of such interventions.

Over the past 20 years, genotyping has informed our knowledge of how TB evolved, spread around the world and survives within hosts(6–8). Unlike some other pathogens, TB genotyping cannot always definitively identify who-infected-whom as multiple cases can be infected with genetically indistinguishable strains (9). Furthermore, cases infected by indistinguishable strains may be epidemiologically unrelated, due to infection with a common strain(10). Instead, genotyping is often used to rule out transmission, for instance between household members infected with different strains(11,12). In low incidence countries, distinguishable unique strains are used to estimate the fraction of cases that are not due to recent transmission, but due instead to the reactivation of existing infections or cases infected elsewhere(13).

TB clusters are defined as multiple cases infected with an indistinguishable genotype. Clusters are often assumed to signify sustained recent transmission. However, their significance depends on the epidemiological setting and may be due to a commonly imported strain. The size of a TB cluster found in a country depends on the number of introductions of a strain, its local transmission potential and the social contact patterns of cases. In the UK, being born in the UK is the biggest risk factor for belonging to a cluster rather than being infected with a unique strain(10). The distribution of cluster sizes has been used to identify critical phenomena in epidemiological processes for acute infections such as measles(14), but is less common for chronic diseases such as TB(15).

Analysis of the distribution of cluster sizes in the Netherlands found that the highly right-skewed distribution of cluster sizes was indicative of super-spreading individuals, i.e. that some individuals generate many more secondary cases than average(16). The method was contingent on identifying transmission clusters (clustered cases occurring less than two years apart): alternative methods are required to apply this method without *a priori* epidemiological knowledge of the likely index case. Furthermore, it is not known how cluster generation differs between settings with potentially differing types of migration and social contact patterns. Here, we propose and develop a method to estimate the average number of secondary cases (the reproduction number) and the percentage of cases that are due to recent transmission from the information in cluster size distributions for TB in the UK and the Netherlands.

## METHODS

### Data sources

#### UK data

The analysis was conducted using TB notifications collected through the Enhanced Tuberculosis Surveillance (ETS) system in England and Wales and the Enhanced Surveillance of Mycobacterial Infections (ESMI) system in Scotland. The following data for TB notifications with isolates with strain typing results of at least 23 loci between 2010 and 2015 were used: year of notification, country of birth, disease type (pulmonary with or without extra-pulmonary or extra-pulmonary only), strain type (24 loci mycobacterial interspersed repetitive unit-variable-number tandem repeat (MIRU-VNTR) type), cluster name (assigned by a PHE naming tool based on strain type) and whether a case was categorised as clustered (yes/no).

#### NL data

Data from NL were extracted from the Netherlands Tuberculosis Register. MIRU-VNTR typing has been systematically conducted in the Netherlands since 2004. As for the UK data, we extracted year of notification (2004-2015), country of birth, disease type (pulmonary or extra-pulmonary), strain type (24 loci mycobacterial interspersed repetitive unit-variable-number tandem repeat (MIRU-VNTR) type).

#### Defining clusters

Cluster size was defined as the number of cases with an indistinguishable MIRU-VNTR profile, where clusters of size 1 were cases with a unique 24 loci VNTR profile. Cases with a single missing locus that matched 23 loci of another cluster were considered part of that cluster(17). Cluster sizes were binned logarithmically to retain the distribution shape while minimising noise due to low numbers of large clusters(18).

#### Mathematical model

We develop a mortal branching process model(16,19) with importation of infection to describe the process by which TB clusters are generated and evolve. The central premise behind the model is that every diagnosed case must have been generated by one of two mechanisms, in a similar structure to household transmission models(15): A) infection was acquired abroad or before the observation period (interpreted as imported infection) or B) the case was infected in the country during the observation period (interpreted as transmitted infection).

For each unique genotype X, we assume that the first case diagnosed with genotype X must have either been infected abroad or before routine genotyping. Each case *i* generates *r*_*i*_ secondary cases infected with genotype X where *r*_*i*_ is drawn from a probability distribution *f* (·). During model development, we considered whether it was necessary to differentiate pulmonary cases from extra-pulmonary cases, due to their differential infectiousness. We found that there was no association between the proportion of pulmonary cases per cluster and cluster size (see results), suggesting the ratio of pulmonary to extra-pulmonary cases is invariant to position in a transmission tree. As extra-pulmonary cases are equally distributed across generations, including extra-pulmonary cases results in a rescaling of the cluster size distribution and will not affect the results. This is consistent will Ypma et al.(16) who found that including extra-pulmonary cases did not affect the results.

In addition to transmission, *m* additional cases infected with genotype X are imported, where *m* is the result of a binomial distribution with probability *p*. During model development, we considered whether importation rate should vary by country of birth. We found that there was no relationship between the proportion of foreign-born cases and cluster size (see results), therefore we modelled importation rate as independent of cluster size.

Each of the generated and imported cases has the opportunity to generate secondary cases; this process is repeated until no new cases are created. The total number of cases infected with genotype X, or the size of a cluster *C*(*X*), is the sum of all the cases:

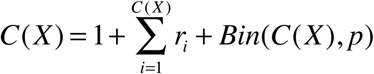

where *r*_*i*_ is the number of secondary cases generated via transmission by case *i* and *Bin*(*C*(*X*), *p*) is the number of imported cases. In order to fit this model to cluster size data without further assumptions, we impose the assumption that the average number of secondary cases per case must be less than one, *E*(*r*_*i*_) < 1, justified by the low and declining incidence in the two countries.

#### Distribution of secondary cases per individual

Ypma et al.(16) modelled the number of secondary cases per TB case using a negative binomial distribution, which arises when the expected number of secondary cases per individual, *λ*, follows a Gamma distribution with dispersion parameter *k* and scaling parameter *θ, λ* ∼ Γ(*k,θ*), where the average number of secondary cases per individual is given by *R* = *kθ*. Initial analysis of the UK data revealed that a negative binomial model was not able to capture the entire distribution of cluster sizes: the best-fit model underestimated the occurrence of both large clusters and single isolates (supplementary figure 1).

We find that a Poisson-lognormal distribution is able to capture the entire distribution of TB cluster sizes for the UK and NL. A Poisson-lognormal distribution is frequently used in ecological literature as an alternative to a negative binomial to describe species abundances for communities with many rare species. It arises when the logarithm of the expected number of secondary cases per individual, log(*λ*), follows a normal distribution with mean *μ* and variance *σ, λ* ∼ log *N* (*μ,σ*). In a lognormal distribution, the average number of secondary cases per individual is given by:

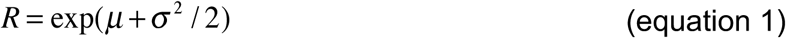

The total number of additional cases due to an average imported case is calculated as:

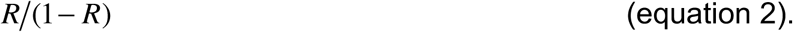

We define a “superspreader” as a TB case in the model that generates more than ten secondary cases. Using the model, we explore the impact of superspreaders by considering the proportion of secondary cases generated by persons with different reproduction numbers.

We use the model to estimate the impact of reducing transmission within the UK to match transmission within the Netherlands. We re-run the model with the estimated UK importation rate but scale the log mean of the Poisson lognormal distribution by a factor 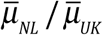, where 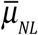 is the average mean of the log normal distribution estimated for the Netherlands and 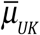 is the average mean of the log normal distribution estimated for the UK. The number of cases is totalled for the alternative scenario with lower transmission and compared to the total number of cases under the UK fitted model.

#### Fraction of imported cases

In the model, a diagnosed case generates an average *R* secondary cases and another case is imported with probability *p*. Therefore, the fraction of imported cases not due to recent transmission is given by *p* (1 + *R* + *p*).

#### Model fitting

In contrast to previous approaches(16,19) that have used exact likelihood methods for fitting cluster size models to data, we use *Approximate Bayesian Computation (ABC)*(20,21). In ABC, the likelihood is approximated by distance metrics based on summary statistics derived from the data and a realisation of the model, therefore can naturally incorporate the impact of sampling and importation. We use the Majoram MCMC search algorithm implemented in the R package EasyABC(22) with the observed frequency of each cluster size as a summary statistic.

From the posterior distributions, we extracted the average number of secondary cases per individual (*R*), the degree of dispersion and the proportion of cases that are due to recent, within-country transmission.

## RESULTS

### Cluster size distributions

In the UK between 2010 and 2015, there were 23,646 genotyped cases and 12,503 unique genotypes. 9,802 cases were unique cases and 13,844 (58.5%) of cases were in clusters containing two or more individuals.

In the Netherlands (NL) between 2004 and 2015, there were 8,449 genotyped cases and 4,955 unique genotypes. 3,905 cases were unique cases and 4,526 (53.6%) of cases were in clusters of two or more. Limiting the analysis to cases diagnosed between 2010 and 2015, there were 3,841 genotyped cases and 2,518 unique genotypes. 2,026 cases had a unique genotype and 1,815 (47·3%) were in clusters of two or more.

### The role of pulmonary cases in cluster formation

In the UK, on average 70% (range 42%, 98%) of clustered cases involved pulmonary disease, compared to 53% of all cases (Figure 1a). In NL, 70% (range 26%, 100%) of clustered cases involved pulmonary disease, compared to 67% of all cases (Figure 1b).

**Figure 1.**
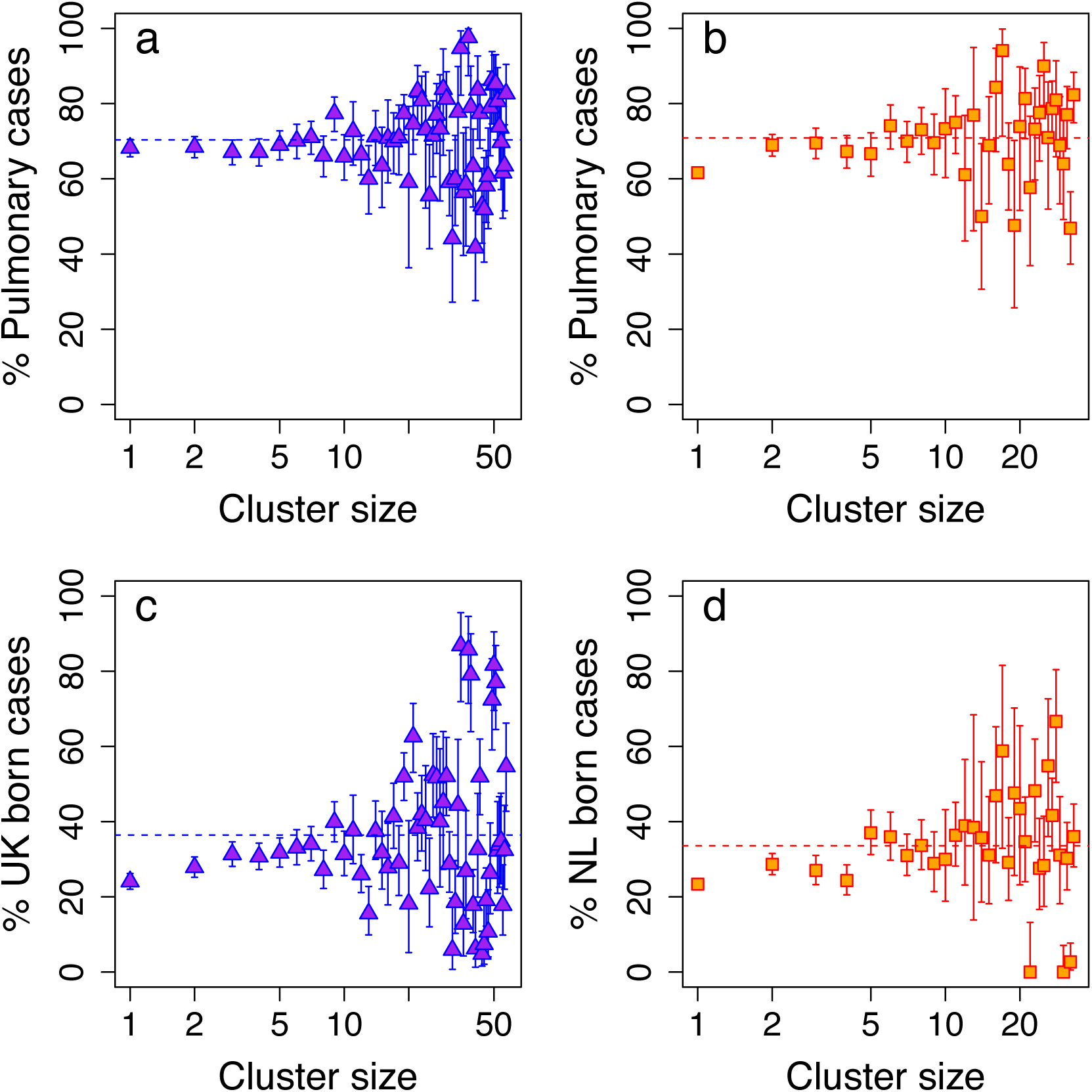
The percentage of pulmonary cases in a cluster against cluster size for the UK (a) and the Netherlands (b). The percentage of foreign-born cases in a cluster against cluster size for the UK (c) and the Netherlands (b). Dotted lines indicate the mean value.

Although having pulmonary disease increased the likelihood of belonging to a cluster, the proportion of cases in a cluster with pulmonary disease had no effect on cluster size in the UK (*R*^2^ = 4 × 10^−4^) or NL (*R*^2^ = 2 × 10^−3^). The invariance of cluster size to the proportion of pulmonary cases suggests that infectious cases generate pulmonary and extra-pulmonary cases and that a similar process operates in both countries.

### The role of foreign-born cases in cluster formation

Foreign-born cases make up a majority, 73% and 69%, of cases in the UK and NL respectively. In both countries, foreign-born cases are less likely to be diagnosed with pulmonary TB (47% vs 69% in the UK and 63% vs 76% in NL) and less likely to belong to a cluster. Foreign-born cases accounted for 64% (range 10%, 95%) of cases in a cluster in the UK and 66% (range 33%, 100%) of cases in a cluster in NL. There was no consistent relationship between cluster size and the percentage of foreign-born cases (p-values>0·1, Figures 1c and 1d).

### Model fit to data

We found that a negative binomial model was unable to capture the extremes of the cluster size distribution in the UK, underestimating either the number of observed unique genotypes or the frequency of large clusters (Supplementary figure 1). Figure 2 shows the Poisson-lognormal model which captures the entire distribution in the UK (a) and NL (b). The Poisson-lognormal distribution for the number of secondary cases in the UK had log-mean of -2·9 (95%CI - 4·7, -1·5) and log-variance 2.0 (95%CI 1·2, 2·8). In NL, between 2004 and 2015 the log-mean was -2·9 (95%CI -5·0, -1·6) and log-variance 1·9 (95%CI 1·0, 2·8). Restricting the analysis to cases reported in NL between 2010 and 2015, decreased the log-mean to -3·4 (95%CI -6·7, -1·7) and slightly increased the log-variance 1·9 (95%CI 1·0, 3·4) (see table S2).

**Figure 2:**
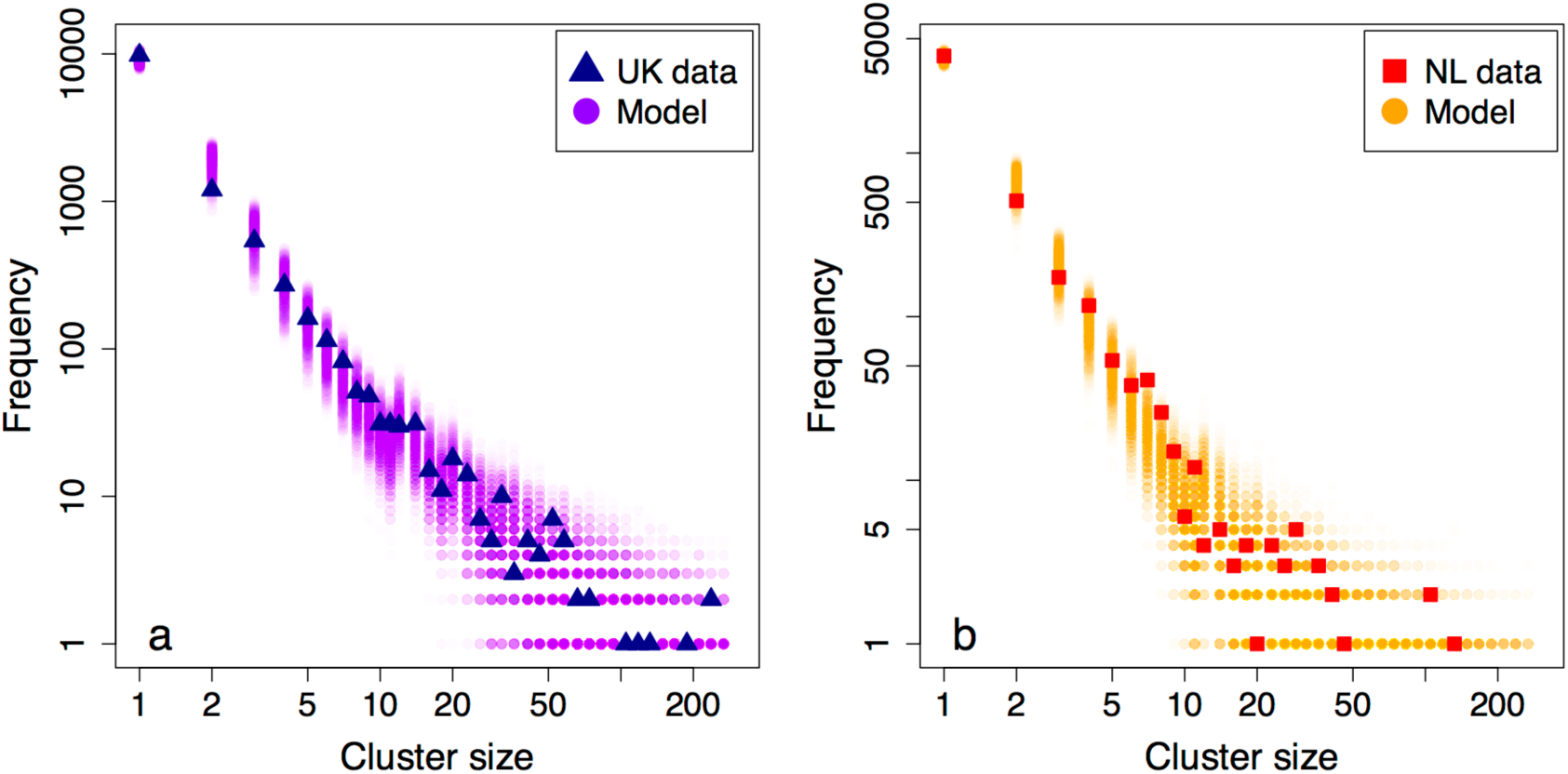
The distribution of cluster sizes for the UK (a) and the Netherlands (b) with the distribution of cluster sizes produced by 1,000 iterations of the Poisson-Lognormal model with parameters drawn from the posterior distributions.

### Cases due to recent transmission

The percentage of imported cases (i.e. not due to transmission in the UK or NL but including cases arising from travelling to country of origin) was estimated at 73% (64%, 79%) in the UK, compared to 81% (69%, 89%) in NL for the same period.

### The Effective Reproduction Number

The mean effective reproduction number is calculated from the Poisson lognormal distribution (equation 1, Methods). For the UK, it was 0·41 (95%CI 0·30, 0·60), suggesting that, on average, transmission is not sustained. Using equation 2 (Methods), this means an average index case will generate 0.7 (0.4, 1.5) further cases. Furthermore, using the model, we find that clusters with more than 10 cases have an average reproduction number greater than 0·9.

In NL, the reproduction number using all data from 2004 to 2015 was 0·33 (95%CI 0·22, 0·50) and since 2010 this reduced to 0·25 (95%CI 0·14, 0·48). Even considering onward transmission chains, an average index case in the Netherlands generates 0.3 (95%CI 0.16, 0.92) further cases.

### The role of superspreaders and onward transmission

Figure 3 illustrates the distribution of secondary cases per diagnosed case. From the UK data, we estimate that 84% (95%CI 73%, 91%) of cases did not generate any secondary cases, therefore current control measures are adequately preventing onward transmission in the majority of cases. A further 10% (95%CI 5%, 18%) of cases generated one secondary case only; they generated 26% (11%, 44%) of cases infected in the UK (figure 3). 0.47% (95%CI 0.2%, 1.2%) of cases generated more than 10 secondary cases, and could be considered “superspreaders”. These superspreaders were responsible for 27% (2%, 55%) of secondary in-country cases, which is less than predicted under a negative binomial model. If one transmission event were prevented for every case, as in a generalised control intervention, 42% (23%, 64%) in-country cases would be prevented.

**Figure 3:**
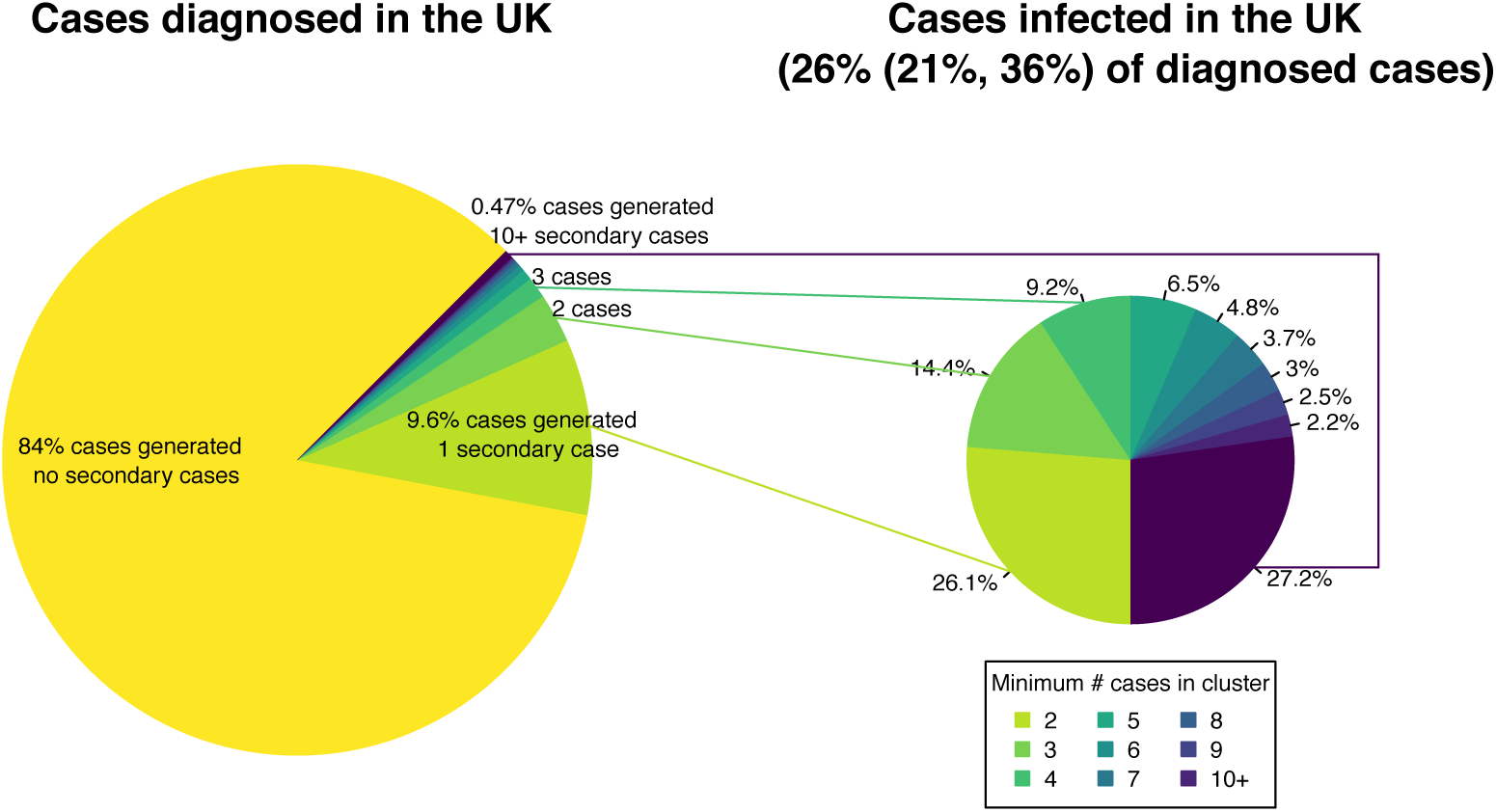
The number of secondary cases produced by diagnosed cases in the UK (left) and those secondary cases as a proportion of cases infected in the UK (right). The point estimates are the mean calculated using 10,000 parameter sets drawn from the posterior distribution of the model fit to the UK data between 2010 and 2015.

In NL between 2010 and 2015, 88% (95%CI 77%, 96%) of cases did not generate any secondary cases and 8% (95%CI 3%, 17%) of cases generated one secondary case, resulting in 34% (9%, 61%) of cases infected in NL. 0.3% (0.02%, 0.8%) of cases were super-spreaders (cases that generate more than 10 secondary cases). These super-spreaders were responsible for 19% (1%, 52%) of recently transmitted cases. Preventing one transmission per case would prevent 53% (20%, 77%) of recently transmitted cases.

By scaling the transmission parameters in the UK, we estimate that if the UK were able to bring local transmission in line with the NL, they would be able to achieve a 17% (13%, 23%) reduction in incidence, equivalent to preventing 538 (266, 818) cases per year.

## DISCUSSION

Tuberculosis (TB) remains a public health concern in low-incidence countries. As the majority of cases in low-incidence countries are foreign-born, impact of controlling recent in-country transmission on overall TB burden is not clear.

Here, we presented methods for using the distribution of TB cluster sizes within a country to characterise on-going transmission. Using data from the UK and NL, we find that the vast majority of cases did not transmit the infection. Less than 1% of cases caused more than 10 secondary cases and could be defined as “superspreaders”. Overall, the average reproduction number is less than a half in both countries.

Superspreading, where a small proportion of cases generate a disproportionate number of secondary cases, is a common feature of many infectious disease epidemics(23). Where superspreading dominates dynamics, targeted interventions perform better than population-wide measures, however identifying superspreaders can be challenging. Our analysis suggests that superspreaders play a less important role than for other diseases, so that control measures applied to all cases are necessary. We estimate that onward transmission is substantially lower in the Netherlands than in the UK. Our estimate of the reproduction number is consistent with previous estimates in low-incidence settings(24). In particular, our estimate is in line with Borgdorff et al.’s estimates that also allow multiple introductions per cluster (25–27), suggesting that this is an important feature. It is lower than Ypma et al.’s estimate, which might be explained by the fact that they allow for the possibility of mutations within a cluster. In the UK, Vynnycky and Fine estimated that the effective reproduction number fell to well below one by 1990 (28). We estimate that the reproduction number is now around 0.4 in the UK, and that reducing this to 0.25, in line with the Netherlands, could prevent one in six UK cases. A comparison of transmission, control measures and outcomes could elucidate the difference we observed between the UK and the NL. These would include the efficiency of contact tracing in the UK(29) and the NL(30), household transmission(12), and different transmission rates between migrant groups(25).

Whole Genome Sequencing (WGS) is increasingly being used for genotyping in high-resource settings(31), having been introduced in 2016 in the NL and 2017 in the UK. Our clustering analysis method could be used to provide an independent estimate of transmission, once the pipeline for TB DNA sequencing has been standardised across countries, consensus reached regarding the cut-off number of SNPs to be used for cluster definition (32) and multiple years of data have accumulated. Even with sufficient data, WGS can be only used to directly estimate reproduction numbers of large outbreaks that extend over a number of years(33), but not outbreaks that occur over short timespans(9). The power of our method lies is the unification of clusters across multiple scales, and is therefore robust to missing data.

However, the approach we used does have limitations. Firstly, the model did not include temporal or regional differences in transmission: these will be areas for future development. Further, we assumed that clusters were fully observed, and that transmission is not sustained without importation from outside the UK/NL or reactivation of old infections. This assumption is a limitation of using a terminal branching process. The steady decline in incidence over the study periods suggests that this assumption is reasonable on average, although transmission is most likely sustained in the largest clusters. However, previous analysis found that right censoring did not affect the overall results(16). We did not capture the role of genetic mutation in generating new clusters, thereby potentially underestimating the contribution of recent transmission. Lastly, our analysis only includes genotyped cases. Including cases that are not culture-confirmed would not alter estimates of transmission, but would impact any cost effectiveness calculations.

In summary, we find that TB clusters in low incidence countries can be explained by the same underlying mechanism of importation and onward transmission and that the mechanism is consistent between countries. The model suggests that super-spreaders are less important than previously thought and that control policies aimed at limiting spread, such as contact tracing, still have a role to play in eliminating TB in low-incidence countries.

## Author contributions

Conceived the study, implemented the analysis: EBP. Developed the R package: LD. Extracted and advised on UK data: JA, ML, CC. Extracted and advised on NL data: HKA, DvS. Interpreted results and wrote manuscript: all.

## Acknowledgements

EBP was supported by the National Institute for Health Research Health Protection Research Unit (NIHR HPRU) in Evaluation of Interventions. The views expressed are those of the author(s) and not necessarily those of the NHS, the NIHR or the Department of Health. The NIHR had no role in writing the manuscript or the decision to publish. EBP had full access to the data and final responsibility for the decision to submit for publication.

## Abbreviations

(TB): Tuberculosis
(UK): United Kingdom
(NL): Netherlands
(WGS): Whole Genome Sequencing
(ETS): Enhanced Tuberculosis Surveillance
(ESMI): Enhanced Surveillance of Mycobacterial Infections
(MIRU-VNTR): Mycobacterial interspersed repetitive unit-variable-number tandem repeat
(PHE): Public Health England

## Data sharing

The cluster size distributions for the UK and NL are in supplementary table 1. The code will be available upon acceptance.

